# Bivalent CD47 Immunotoxin for Targeted Therapy of Lung Cancer

**DOI:** 10.64898/2025.12.24.696420

**Authors:** Jihong Ma, Dinoop Ravindran Menon, Zhaohui Wang, Danielle Mintzlaff, Hugh Rock, Lauren Giesy, Christene A. Huang, David W. Mathes, Trevor Nydam, Elizabeth A. Pomfret, D. Ross Camidge, Hatim E. Sabaawy, Sharon R. Pine, Zhirui Wang

**Author notes:** Corresponding Authors: Zhirui Wang, PhD, Department of Surgery, School of Medicine, University of Colorado Anschutz Medical Campus, RC2, Rm 6013, Mail Stop: 8621, 12700 E 19^th^ Ave., Aurora, CO 80045, USA, Sharon R. Pine, PhD, Division of Medical Oncology, Department of Medicine, University of Colorado School of Medicine, 12801 East 17th Ave, Mail Stop: 8117, Aurora, CO 80045, USA.

## Abstract

Lung cancer is the leading cause of cancer-related deaths in the world, with ∼2.5 million people diagnosed and ∼1.5 million deaths each year. While the last two decades have yielded substantial progress with systemic targeted and immune therapies improving treatment responses in subgroups of patients with advanced and refractory lung cancer, there is still a dire unmet clinical need to develop more effective therapies with durable responses. CD47 receptors are overexpressed on the surface of a variety of malignant tumor cells including lung cancer. Although there has been increasing interest in developing targeting antibodies against CD47 for immunotherapy, they failed clinical trials due to their dose-limiting toxicities, primarily hematopoietic toxicity. Using a unique diphtheria toxin resistant *Pichia pastoris* yeast expression system, we have developed a diphtheria toxin-based bivalent CD47 immunotoxin (bi-CD47-IT) for targeted therapy of CD47^+^ non-small cell lung cancer (NSCLC). Bi-CD47-IT demonstrated compelling preclinical efficacy in multiple NSCLC cell line-derived xenograft (CDX) and patient-derived xenograft (PDX) mouse models, including subcutaneous, orthotopic, metastatic and humanized models. This study demonstrates the remarkable preclinical activity of bi-CD47-IT against various lung cancer models, making bi-CD47-IT a novel and promising therapeutic approach for NSCLC.

## Introduction

Lung cancer remains the leading cause of cancer-related mortality worldwide, with nearly 2.5 million new cases reported and over 1.5 million deaths annually (1). The overall 5-year survival rate for advanced non-small lung cancer (NSCLC) varies from 9–12%. Improvements in systemic therapy including antibodies, cytotoxic chemotherapy, and especially tyrosine kinase inhibitors (TKIs) have modestly extended overall survival. However, the vast majority of patients treated with systemic targeted therapies will experience progression, and only about 20% of advanced NSCLC patients treated with immunotherapy plus chemotherapy survive up to 5 years (2,3). Consequently, prognosis for advanced NSCLC remains dismal.

The CD47/SIRPα (CD172a) interaction is one of the most well-characterized immune checkpoints that control macrophages in tumors (4). CD47 is highly expressed on many cancers including lung cancer, and it binds to SIRPα, a receptor expressed by macrophages and other myeloid cells. When CD47 and SIRPα engage, inhibitory signals are activated by macrophages to prevent phagocytosis. Expression of CD47 in cancer cells can be controlled by cytokines, oncogenes, and micro-RNAs (5). Studies have shown that EGFR-mutant tumors are enriched for high CD47 expression at both the mRNA and protein levels (6–8). Recent studies established oncogenic mechanisms of CD47 expression regulation by lung cancer oncogenic drivers, EGFR and KRAS (9,10). In addition to oncogenes and cytokines, clinical studies demonstrated that standard of care lung cancer therapies upregulate CD47 expression, many through the JAK/STAT pathway (11,12). CD47 in tumor cells demonstrated interactions with proteins that collectively influence cell death and survival, migration, invasion, and angiogenesis (13). High CD47 expression in NSCLC was shown in large cohorts of patients to be associated with advanced tumor stage and poor patient survival (6,7,14).

The CD47-SIRPα axis is a highly recognized target for developing blocking monoclonal antibodies and recombinant fusion proteins for cancer immunotherapy. Previous approaches were designed to block the interaction between CD47 and SIRPα to relieve suppression of cancer cell phagocytosis by antigen presenting cells in order to stimulate tumor immunity. However, monotherapy with previous CD47 blockade was ineffective in human clinical trials against many tumor types (15). Moreover, both CD47 mAbs and SIRPα–Fc fusion proteins have generally been well tolerated in patients, but grade 3–4 toxicities due to CD47 blockade on RBCs, platelets, and neutrophils occurred in 18% of patients (16), with the most common being thrombocytopenia, neutropenia, and anemia, consistent with high levels of CD47 expression on platelets, neutrophils, and RBCs, respectively. Fc-FcγR interactions are required for the antitumor activities of anti-CD47 antibodies (17). Using a unique diphtheria toxin-resistant *Pichia pastoris* yeast expression system (18), we have developed a truncated diphtheria toxin DT390-based bivalent anti-human CD47 immunotoxin (bi-CD47-IT) for targeted therapy of CD47^+^ cancers utilizing the immunotoxin (IT) approach (19). We genetically engineered bi-CD47-IT to contain two-tandem anti-human CD47 scFv moieties fused to a truncated diphtheria toxin, DT390 (20). In a prior study, we demonstrated that bi-CD47-IT had compelling *in vivo* efficacy against T-cell acute lymphoblastic leukemia (T-ALL) with very low toxicity due to the low binding avidity of bi-CD47-IT to CD47. Bi-CD47-IT significantly prolonged the median survival of T-ALL tumor-bearing mice and highly effectively depleted the T-ALL blast cells in the peripheral blood, spleen, liver, bone marrow, brain, and spinal cord in all tested early and overt T-ALL CDX and PDX mouse models (20). Because CD47 is highly expressed in lung cancer, herein we have tested the anti-tumor efficacy of bi-CD47-IT in lung tumors.

## Materials and Methods

### Human NSCLC cell lines, human NSCLC PDX samples, bivalent CD47 immunotoxin, and antibodies

Human NSCLC A549 and H1299 cell lines were obtained from ATCC (Cat# CCL-185 and CRL-5803, Manassas, VA). Human NSCLC PDX1 (CT257-F4) was obtained from Apar Pataer (MD Anderson Cancer Center) and PDX2 (220900-M3) was generated from an EML4-ALK fusion-positive patient who progressed on ALK-targeted therapy at the University of Colorado. Flow cytometry analysis confirmed that the two NSCLC PDX samples were CD47^+^ (Supplementary Fig. S6E-F). The NSCLC PDX tumors were expanded in immunodeficient *NSG* mice. When the NSCLC PDX tumor volume reached the maximum allowed volume, the tumor-bearing mice were euthanized and the PDX tumors were harvested, cut to small pieces, and directly implanted into the dorsal flank of the experimental *NSG* mice. Bivalent CD47 immunotoxin (bi-CD47-IT) was produced in our laboratory (20) The antibodies used in this study are listed in Supplemental Table 1.

### Alexa Fluor 488-labeling of bi-CD47-IT

Bi-CD47-IT was labeled using Alexa Fluor 488 microscale protein labeling kit (Thermo Fisher Scientific) following the manufacturer’s instruction. In brief, bi-CD47-IT was concentrated down to a concentration of ∼1 mg/mL, The 1/10 volume of sodium bicarbonate solution was added to the protein solution and mixed well. An appropriate volume of Alexa Fluor 488 reactive dye solution was added according to the equation of the manufacturer’s instruction and incubated for 15 minutes at room temperature. The conjugate reaction mixture was uploaded onto the resin bed surface and centrifuged at 16,000× g for 1 minute. The purified dye-labeled bi-CD47-IT was collected into an Eppendorf tube. The labeled-immunotoxin concentration was measured using nanodrop and stored at 4℃ for use.

### Flow cytometry binding avidity analysis of bi-CD47-IT

Human CD47^+^ NSCLC cells (A549 and H1299) were stained with Alexa Fluor 488-labeled bi-CD47-IT at a range of concentrations (1 to 1000 nM). The FITC-labeled anti-human CD47 mAb (BD Bioscience, Cat# 556045, San Jose, CA) at 200 nM was used as a positive control. Alexa Fluor 488-labeled isotype mouse IgG1 served as a negative control at a final concentration of 200 nM. The 1×10^6^ tumor cells were aliquoted into a 12×75 mm tube and incubated with serial diluted Alexa Fluor 488-labeled bi-CD47-IT for 30 minutes at 4℃. The tumor cells were washed with 2 mL of cold flow cytometry buffer twice and spun down at 300× g for 5 minutes. The supernatant was discarded and resuspended in 300 µL of flow cytometry buffer. 10 µL of 7-AAD was added and stained at room temperature for 10 minutes. Flow cytometry was carried out using a CytoFLEX Flow cytometer (Beckman Coulter, Brea, CA), and the data was analyzed using FlowJo software (FLOWJO, LLC, Ashland, OR).

### K_D_ determination

Binding of the bi-CD47-IT to human CD47^+^ NSCLC cells was performed using a wide concentration range (1-1000 nM) of Alexa Fluor 488-labeled bi-CD47-IT. K_D_ determination was performed based on the flow cytometry data using nonlinear regression with the saturation binding equation (GraphPad Prism 10, San Diego, CA). The median fluorescence intensity (MFI) was plotted versus the Alexa Fluor 488-labeled bi-CD47-IT concentrations. Nonlinear regression was based on the equation Y = Bmax × X/(K_D_ + X), where Y = MFI at the given Alexa Fluor 488-labeled bi-CD47-IT after subtracting the background, X = Alexa Fluor 488-labeled bi-CD47-IT concentration, and Bmax = the maximum specific binding in the same units as Y. The same procedure was applied for the flow cytometry binding affinity analysis and K_D_ determination of human red blood cells, human lymphocytes, and human monocytes.

### In vitro efficacy analysis of bi-CD47-IT

The *in vitro* efficacy of bi-CD47-IT was determined in human CD47^+^ NSCLC cells using the CellTiter-Glo^®^ Luminescent Cell Viability Assay (Promega, Madison, WI) as described previously (21). This assay measures the luminescence produced by ATP production from metabolically active cells. Increasing concentrations of bi-CD47-IT cause cell death and a corresponding reduction in ATP-related fluorescence. The luminescence signals were recorded using a BioTek Synergy LX Multi-Mode Reader (Agilent, Santa Clara, CA). The pCD3-IT (anti-porcine CD3 immunotoxin) was included as negative immunotoxin control. In brief, 1×10^4^ tumor cells in 100 µL were added to each well of the 96-well plate. The serial diluted immunotoxins starting at 1×10^-7^ M to 1×10^-14^ M were added and mixed well to a triple-well set for each diluted immunotoxin. The reaction plate was incubated for 48 hours at 37℃ with 5% CO_2_. The plate was equilibrated at room temperature for 30 minutes. An equal volume of the pre-prepared lyophilized enzyme/substrate CellTiter-Glo mixture was added to each well. The plate was mixed for 2 minutes at 1000 rpm on a shaker to induce cell lysis. The plate was incubated at room temperature for 10 minutes to stabilize luminescent signal in the dark. The luminescence signal was measured using the microplate reader.

### In vivo efficacy studies of bi-CD47-IT

All *in vivo* experiments were approved by the University of Colorado Anschutz Medical Campus Animal Care and Use Committee. The lung tumor volume was measured using digital vernier calipers and calculated according to the formula: volume (mm^3^) = [length × (width × 2)]/2 (22). Mice were euthanized at their end point defined as when the tumor volume became more than 1000 mm^3^, or body weight loss was more than 15%. Bi-CD47-IT was assessed in the following mouse models. **1) Human NSCLC A549 CDX *NSG* mouse model**. *NSG* mice (7-8 weeks old from The Jackson laboratory) were used for this study. Human NSCLC A549 cells (5 million) were subcutaneously injected into the dorsal flank of the *NSG* mice. When the tumor volume reached ∼50 mm^3^ (day 4 post the A549 tumor cell injection), the tumor-bearing mice were treated with 81 µg/kg of bi-CD47-IT by daily intraperitoneally injection for 10 consecutive days. The dose schedule of bi-CD47-IT was based on our previous experience (20). PBS was included as a vehicle treatment control. The tumor volume was monitored twice weekly, and the tumor volume curve was recorded. When the first tumor in PBS group reached the maximum allowed volume (∼1000 mm^3^) on day 40 post the A549 tumor cell injection, all tumor-bearing mice were euthanized to collect tumors for comparison of the tumor size and weight. **2) Human NSCLC H1299 CDX *NSG* mouse model**. *NSG* mice (7-8 weeks old from The Jackson laboratory) were used for this study. Human NSCLC H1299 cells (5 million) were subcutaneously injected into the dorsal flank of the *NSG* mice. When the tumor volume reached ∼70-80 mm^3^ (day 4 post the H1299 tumor cell injection), the tumor-bearing mice were treated with 81 µg/kg of bi-CD47-IT by daily intraperitoneally injection for 10 consecutive days. PBS was included as a vehicle treatment control. The tumor volume was monitored twice weekly, and the tumor volume curve was recorded. When the first tumor in PBS group reached the maximum allowed volume (day 28 post the H1299 tumor cell injection), all tumor-bearing mice were euthanized to collect the tumors for comparison of the tumor size and weight. **3) Human NSCLC A549 CDX humanized mouse model**. Hu-CD34^+^ NSG-SGM3 mice (14 weeks old) were used for this study. Human NSCLC A549 cells (5 million) were subcutaneously injected into the dorsal flank of the humanized mice. The chimerism level was confirmed by flow cytometry analysis (>25%). When the tumor volume reached ∼70 mm^3^ (day 5 post the A549 tumor cell injection), the tumor-bearing mice were treated with 81 µg/kg of bi-CD47-IT by daily intraperitoneal injection for 10 consecutive days. PBS treatment group was included as vehicle treatment control. Tumor volumes were monitored twice weekly. When the first tumor in the PBS treatment group reached the maximum allowed volume on day 33, all tumor-bearing mice were euthanized to collect the tumors for comparison of the tumor size and weight. **4) Orthotopic human NSCLC A549 CDX *NSG* mouse model**. *NSG* mice (7-8 weeks old from The Jackson laboratory) were used for this study. Human NSCLC A549 cells (500,000) were orthotopically injected into the left lung of the *NSG* mice. When the orthotopic A549 NSCLC was detectable by micro-CT scan analysis (day 10 post the tumor cell injection), the tumor-bearing mice were treated with 81 µg/kg of bi-CD47-IT by daily intraperitoneal injection for 10 consecutive days. PBS treatment group was included as vehicle treatment control. The survival curve was recorded. The tumor volume was monitored once weekly by micro-CT scan analysis. When the first tumor-bearing mouse in the PBS treatment group met the study endpoint on day 25 post the A549 tumor cell injection, two tumor-bearing mice were euthanized for gross examination and histological analysis. The remaining tumor-bearing mice were continually monitored to the study endpoint. **5) Orthotopic human NSCLC A549 CDX humanized mouse model**. Hu-CD34^+^ NSG-SGM3 mice (12 weeks old) were used for this study. The chimerism level was confirmed by flow cytometry analysis (>25%). Human NSCLC A549 cells (500,000) were orthotopically injected into the left lung of Hu-CD34^+^ NSG-SGM3 mice. When the lung tumor was detectable by micro-CT scan analysis on day 7 post the A549 tumor cell injection, the tumor-bearing mice were treated with 81 µg/kg of bi-CD47-IT by daily intraperitoneal injection for 10 consecutive days. PBS treatment group was included as vehicle treatment control. The survival curve was recorded. Lung tumor volume was monitored once weekly by micro-CT scan analysis. When the first tumor-bearing mouse in the PBS treatment group met the study endpoint on day 29 post the A549 tumor cell injection, two tumor-bearing mice were euthanized for gross examination and histological analysis. The remaining tumor-bearing mice were continually monitored to the study endpoint. **6) Experimental metastasis human NSCLC A549 CDX *NSG* mouse model**. *NSG* mice (7-8 weeks old from The Jackson laboratory) were used for this study. Human NSCLC A549 cells (2 million) were intravenously injected into the *NSG* mice via tail vein. Bi-CD47-IT treatment started on day 4 at 81 µg/kg, intraperitoneal injection, once daily for 10 consecutive days. PBS treatment group was included as vehicle treatment control. The survival curve was recorded. When the first tumor-bearing mouse in the PBS treatment group met the study endpoint on day 38 post the A549 tumor cell injection, two tumor-bearing mice were euthanized for gross examination and histology analyses of lungs, spleens, livers and kidneys. The remaining tumor-bearing mice were continually monitored to the study endpoint. **7) Human NSCLC PDX1 (CT257-F4) *NSG* mouse model**. *NSG* mice (7-8 weeks old from The Jackson Laboratory) were used for this study. NSCLC PDX1 tumor sample was amplified using *NSG* mice. The amplified fresh NSCLC PDX1 tumors were cut into small pieces and subcutaneously implanted into the dorsal flank of the experimental *NSG* mice. When the tumor was palpable on day 15 post the PDX1 tumor implantation, bi-CD47-IT treatment started at 81 µg/kg, intraperitoneal injection, once daily for 10 consecutive days. PBS treatment group was included as vehicle treatment control. Tumor volumes were monitored on day 15, 25 and 44. When the first tumor in the PBS treatment group reached the maximum allowed volume on day 44 post the PDX1 tumor implantation, all tumor-bearing mice were euthanized to collect the tumors for comparison of the tumor size and weight. **8) Human NSCLC PDX2 (220900-M3) *NSG* mouse model**. *NSG* mice (7-8 weeks old from The Jackson Laboratory) were used for this study. Human NSCLC PDX2 tumor sample was amplified using *NSG* mice. The amplified fresh NSCLC PDX2 tumors were cut into small pieces and subcutaneously implanted into the dorsal flank of the experimental *NSG* mice. When the tumor was palpable on day 12 post the PDX2 tumor implantation, bi-CD47-IT treatment started at 81 µg/kg, intraperitoneal injection, once daily for 10 consecutive days. PBS treatment group was included as vehicle treatment control. Tumor volumes were monitored on days 12, 22, 30 and 35. When the first tumor in the PBS treatment group reached the maximum allowed volume (day 35), all tumor-bearing mice were euthanized to collect tumors for comparison of the tumor size and weight. **9) *In vivo* efficacy comparison of bi-CD47-IT versus Cisplatin in a human NSCLC A549 CDX *NRG* mouse model.** *NRG* mice (7-8 weeks old from The Jackson laboratory) were used for this study. NSCLC A549 tumor cells (5 million) were subcutaneously injected into the dorsal flank of the *NRG* mice. When the tumor volume reached ∼50 mm^3^ (day 4 post the tumor cell injection), bi-CD47-IT treatment started at 81 μg/kg, intraperitoneal injection, once daily for 10 consecutive days. Cisplatin was intraperitoneally injected at 5 mg/kg (23,24), once weekly for 2 doses total. The tumors were monitored twice weekly, and the tumor volume curve was recorded. When the first tumor in the PBS treatment group reached the maximum allowed volume (day 28), all tumor-bearing mice were euthanized to collect the tumors for comparison of the tumor size and weight.

### Sex as a biological variable

Same age and equal numbers of male and female *NSG* mice from the Jackson Laboratory were used for each experiment. Only female humanized Hu-CD34^+^ NSG-SGM3 mice were used in this study because human chimerism (the presence of human cells within a mouse’s system) is generally more stable in female humanized mice than in male humanized mice.

### Histology analysis

Murine tissues were surgically harvested from the tumor-bearing mice. The collected tissues were fixed in 10% formalin, embedded in paraffin, and subsequently sectioned. Hematoxylin and eosin (H&E) staining was performed by the University of Colorado Histology Shared Resource Core. CD47 expression was analyzed by immunohistochemistry (IHC) staining using an anti-human CD47 mAb (40 µg/mL, Invitrogen), followed by a goat anti-mouse HRP-conjugated secondary antibody. Ki-67 expression was analyzed by IHC staining using an anti-Ki-67 mAb (Biocare Medical) according to the manufacturer’s protocol. All slides were incubated with the primary antibodies overnight at 4°C, followed by incubation with the secondary antibodies for 1 hour at room temperature. Nuclear staining was performed with hematoxylin for IHC. Images were captured using an Echo Revolve Microscope (San Diego, CA).

### Statistical Analysis

The *p*-values for the survival curves were calculated using the Mantel-Cox log-rank test (GraphPad Prism 10). The *p*-values for the tumor growth inhibition curves were calculated using Two-way ANOVA (GraphPad Prism 10). The *p*-values for other comparisons were calculated using the two-tailed Student t-test (GraphPad Prism 10). *P* < 0.05 was considered statistically significant.

## Results

### Bi-CD47-IT effectively binds to and inhibits human non-small cell lung cancer (NSCLC) cells

We first evaluated the *in vitro* binding activity of Alexa Fluor 488-labeled bi-CD47-IT to two representative human NSCLC cell lines by flow cytometry. We chose A549 cells with an activating KRAS G12S mutation, and inactivating STK11, KEAP1 and CDKN2A genomic changes, and H1299 cells with activating PIK3CA E545K and NRAS Q61K mutations and loss of TP53 and CDKN2A. We confirmed that both cell lines had high expression of CD47. A549 and H1299 cells express 9.7-fold and 8.2-fold more CD47, respectively, than human red blood cells (Supplementary Fig. S1A). As shown in **Fig. 1A and 1D**, bi-CD47-IT bound to A549 and H1299 cells in a dose-dependent fashion with K_D_ values of 916.5 nM and 5,829 nM, respectively (**Fig. 1B and 1E**). We next tested the efficacy of bi-CD47-IT against these cell lines *in vitro* using a CellTiter-Glo^®^ luminescent cell viability assay. Bivalent anti-porcine species-specific CD3 immunotoxin (pCD3-IT) was used as a negative immunotoxin control in mice. The pCD3-IT was genetically engineered to link the same truncated diphtheria toxin, DT390, with bivalent anti-porcine CD3 epsilon scFvs (25). Bi-CD47-IT very effectively inhibited the growth of A549 cells, with an IC_50_ value of 1.58 × 10^-11^ M (**Fig. 1C**), and inhibited H1299 cells, with an IC_50_ of 0.97 × 10^-10^ M (**Fig. 1F**). In contrast, pCD3-IT had no effect on cell viability. These data demonstrate that bi-CD47-IT is highly potent against NSCLC *in vitro*, despite its low binding affinity.

**Figure 1.**
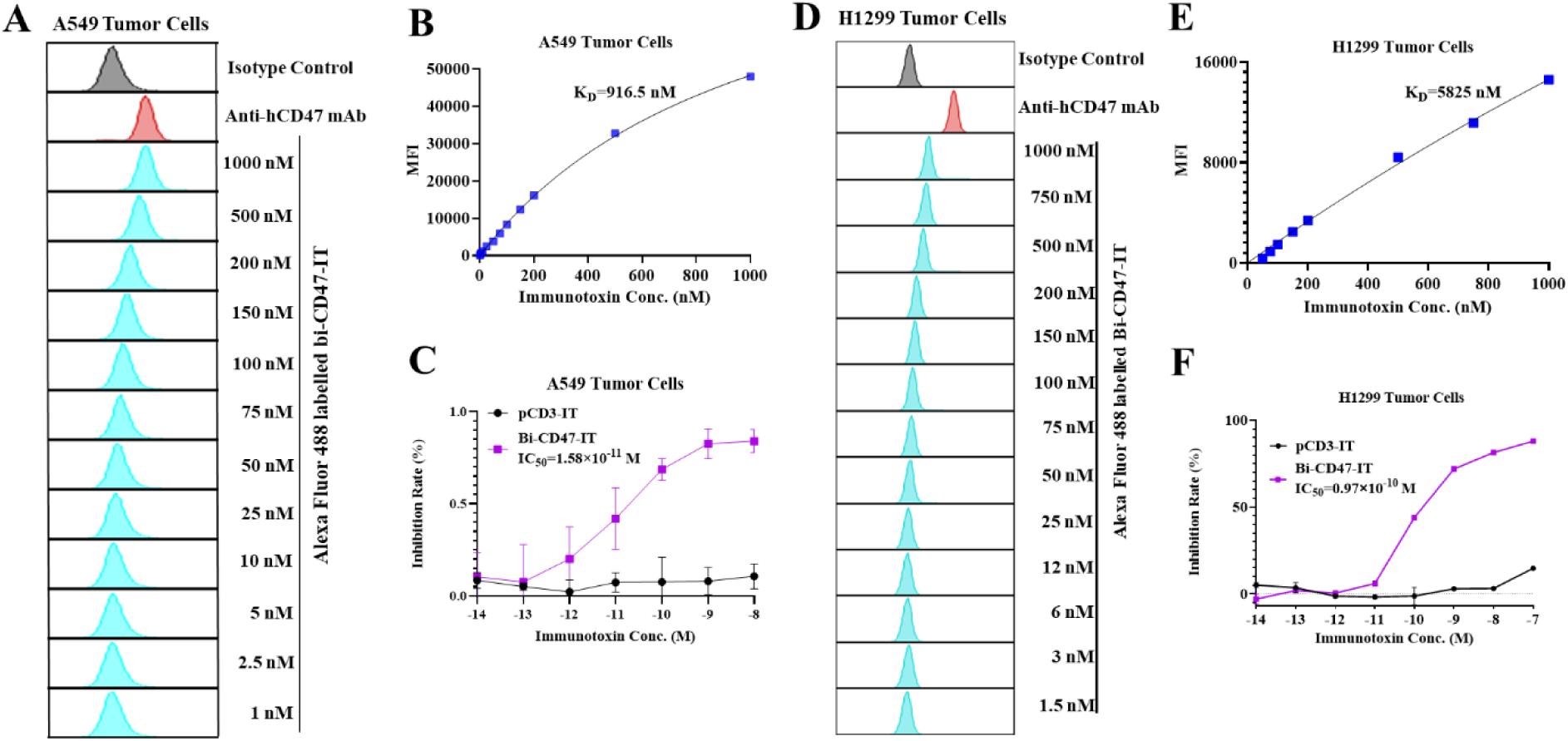
Bi-CD47-IT bound and inhibited two non-small cell lung cancer (NSCLC) cells. **(A and D)** *In vitro* binding affinity analysis of the Alexa Fluor 488-labeled bi-CD47-IT to human NSCLC A549 cells **(A)** and human NSCLC H1299 cells **(D)** by flow cytometry analysis. FITC-anti-human CD47 mAb (B6H12) was used as a positive control, and Alexa Fluor 488-labeled isotype mouse IgG1 served as a negative control. The data are representative of three individual experiments. **(B and E)** K_D_ determination of bi-CD47-IT to human NSCLC A549 cells **(B)** and human NSCLC H1299 cells **(E)** using flow cytometry and nonlinear least-squares fitting. The mean fluorescence intensity (MFI) was plotted over a wide range of Alexa Fluor 488-labeled bi-CD47-IT concentrations. Nonlinear regression was based on the equation Y = Bmax × X/ (K_D_ + X), where Y = MFI at the given Alexa Fluor 488-labeled bi-CD47-IT after subtracting the background, X = Alexa Fluor 488-labeled bi-CD47-IT concentration, and Bmax = the maximum specific binding in the same units as Y. **(C and F)** *In vitro* efficacy of bi-CD47-IT to human NSCLC A549 cells **(C)** and human NSCLC H1299 cells **(F)** determined by the CellTiter-Glo^®^ Luminescent Cell Viability Assay (purple line: bi-CD47-IT group; black line: pCD3-IT group as negative control). Y-axis: percent inhibition of cell viability determined by the number of viable cells based on the quantification of ATP. X-axis: immunotoxin concentration. Cycloheximide (1.25 mg/mL) was used as a positive control. The negative control wells contained cells without immunotoxin. Data is from three individual experiments. Error bars indicate SD.

### Bi-CD47-IT effectively inhibits tumor growth in two NSCLC CDX mouse models

We next explored the *in vivo* efficacy of bi-CD47-IT against NSCLC. A549 cells were subcutaneously injected into the dorsal flank of *NSG* (NOD scid gamma mice) mice, and once the average tumor volume reached ∼50 mm^3^ (day 4 post tumor cell injection), mice were treated with 81 µg/kg of bi-CD47-IT by daily intraperitoneal injections for 10 consecutive days (Supplementary Fig. S2A). The dosing schedule was based on our previous experience (20). As shown in **Fig. 2A-D**, compared with vehicle control, bi-CD47-IT showed significant anti-tumor activity (*P*<0.0001), as measured by tumor volume and tumor weight. There was a significant reduction in tumor volume as early as 10 days after the start of treatment, in which 4 of the 6 tumors were no longer palpable (**Fig. 2B**), and tumors were still not detectable in 50% (3 of 6) of the mice at the end of the study (**Fig. 2C**). We next validated these findings in an H1299 subcutaneous CDX mouse model (Supplementary Fig. S2C). As shown in **Fig. 2E-H**, bi-CD47-IT also significantly inhibited *in vivo* tumor growth in these cells (*P*<0.0001). Thus, bi-CD47-IT has potent anti-tumor activity against NSCLC *in vitro* and *in vivo*.

**Figure 2.**
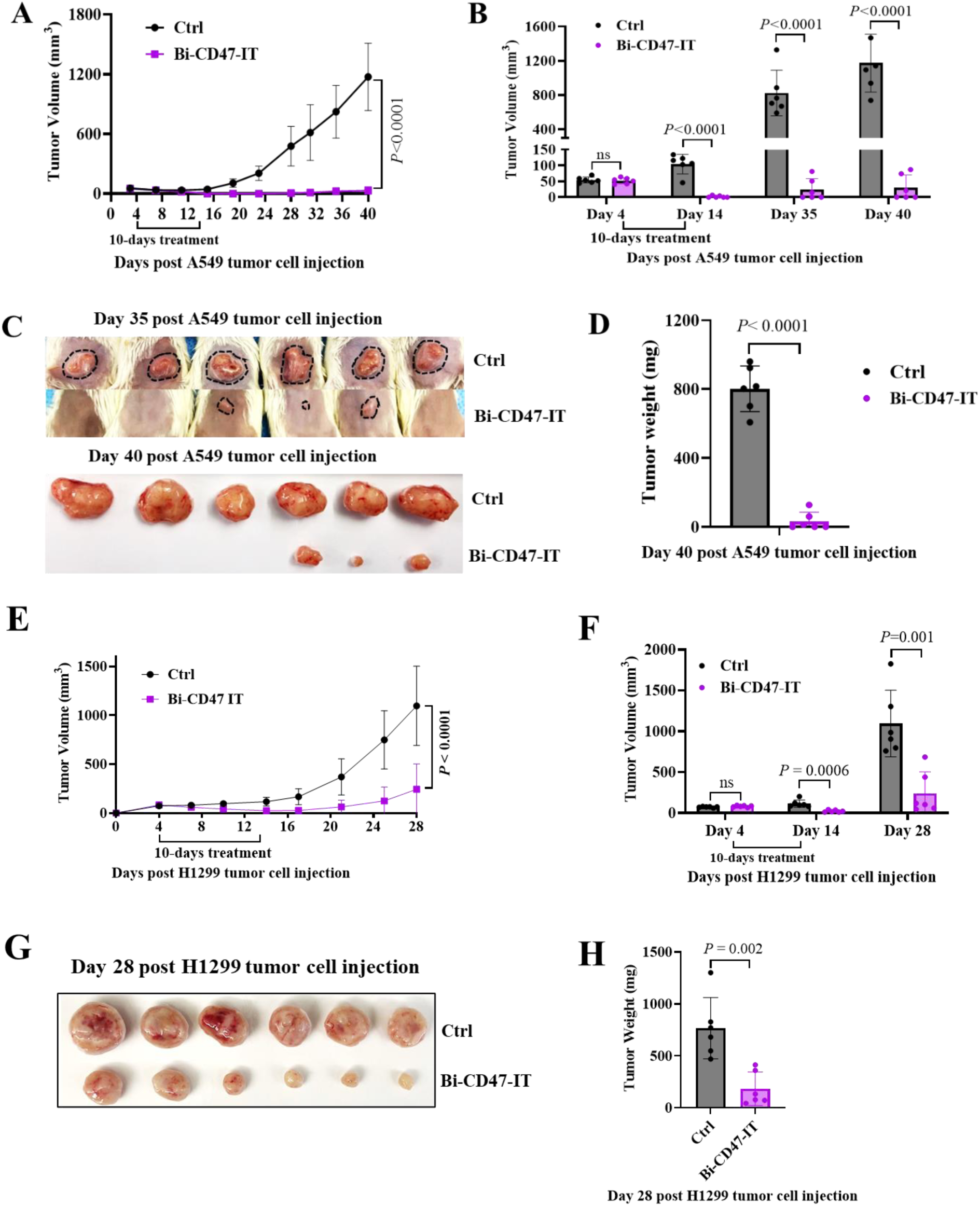
Bi-CD47-IT effectively inhibited tumor growth in two NSCLC CDX mouse models. **(A-D)** Human NSCLC A549 CDX mouse model (*NSG* mice). **(A)** A549 tumor volume curves. **(B)** A549 tumor volume on day 4 (pre-treatment), 14 (post-treatment), 35 and 40 (study endpoint). **(C)** A549 tumor volume on day 35 and tumor size on day 40. **(D)** A549 tumor-weight on day 40. No visible tumors were found on day 40 in 50% (3 of 6) of the tumor-bearing mice. **(E-H)** Human NSCLC H1299 CDX mouse model (*NSG* mice). **(E)** H1299 tumor volume curves. **(F)** H1299 tumor volume on day 4 (pre-treatment), 14 (post-treatment), and 28 (study endpoint). **(G)** H1299 tumor size on day 28. **(H)** H1299 tumor weight on day 28. The *p* values in panel **(A)** and **(E)** were calculated using Two-Way ANOVA (GraphPad Prism 10). The *p* values in panel **(B), (D), (F)** and **(H)** were calculated using two-tailed Student t-test (GraphPad Prism 10).

### Bi-CD47-IT demonstrates antitumor activity in humanized and orthotopic NSCLC A549 CDX mouse models

Immunodeficient *NSG* mouse models have their limitation for the efficacy study, which limits the assessment of the possible interaction between bi-CD47-IT treatment versus the tumor microenvironments including human immune cells (26). We next assessed the *in vivo* efficacy of bi-CD47-IT in the presence of human immune system, using a humanized mouse model (Hu-CD34^+^-NSG-SGM3 mice) (Supplementary Fig. S3A). As shown in **Fig. 3A-D**, bi-CD47-IT significantly inhibited A549 CDX tumor growth in the presence of the human immune system (*P*<0.0001). To mimic the advanced stages of the disease, we started the treatment later when the tumors were slightly larger for the bi-CD47-IT treatment in the humanized mice as compared to the *NSG* mice (∼70 mm^3^ vs ∼50 mm^3^), and there was still substantial anti-tumor activity. No weight loss was observed, which indicates good safety of bi-CD47-IT treatment in the humanized mouse model (Supplementary Fig. S3B).

**Figure 3.**
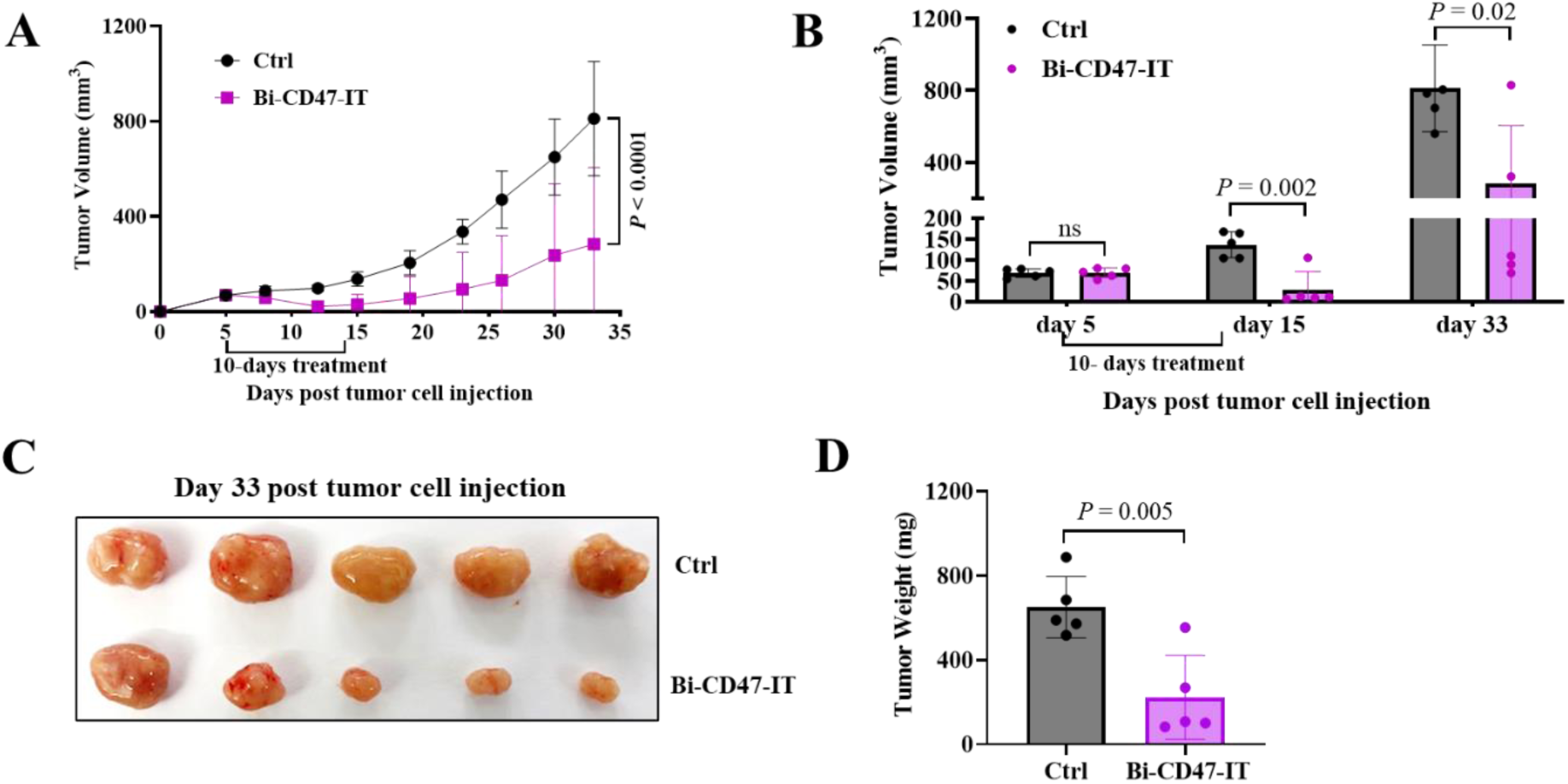
Bi-CD47-IT effectively inhibited tumor growth in a humanized NSCLC A549 CDX mouse model. Hu-CD34^+^ NSG-SGM3 mice were used for this study. **(A)** Tumor volume curves. **(B)** Tumor volume on day 5 (pre-treatment), 15 (post-treatment), and 33 (study endpoint). **(C)** Tumor size on day 33. **(D)** Tumor weight on day 33. The *p* value in panel **(A)** was calculated using Two-Way ANOVA (GraphPad Prism 10). The *p* values in panel **(B)** and **(D)** were calculated using two-tailed Student t-test (GraphPad Prism 10).

Because the tumor microenvironment is known to affect anti-tumor activity and orthotopic models have the potential to increase the predictive values of preclinical studies on patient response (27) (28), we next tested the efficacy of bi-CD47-IT in an orthotopic model (Supplementary Fig. S4A). We injected A549 cells (500,000) into the left lung of *NSG* mice, and when the lung tumors were detectable by micro-CT scan analysis (day 10 after tumor cell injection), the mice were treated with bi-CD47-IT as detailed above. As shown in **Fig. 4D**, bi-CD47-IT significantly prolonged the survival of the tumor-bearing mice from a median survival of 25 days in the vehicle control-treated mice to 40 days in the bi-CD47-IT-treated mice. When the first tumor-bearing mouse in the control group reached the study endpoint (day 25 post the tumor cell injection), two random mice were euthanized from each group. Widespread tumor nodules were observed in the entire lungs as well as the chests, diaphragms and abdomens of the controls (**Fig. 4E**), as expected based on our prior experience with this cell line (data not shown). In contrast, no or less tumor nodules in the lungs and no metastatic lesions were observed in mice treated with bi-CD47-IT. Histology analysis (H&E staining) data was consistent with the survival curve and gross examination data. Large tumor masses were observed in the lungs of the control mice, and only minimal or no residual tumors were observed in the lungs of bi-CD47-IT-treated mice, while regaining pre-implantation lung architecture (**Fig. 4F**). The micro-CT scan image analysis data was also consistent with the survival curve, gross examination and histology analysis (H&E staining) data, with a significant reduction in tumor size in the bi-CD47-IT-treated mice (compared to the control group, *p*<0.001) (**Fig. 4A-C**). We further confirmed the efficacy of bi-CD47-IT in a humanized orthotopic NSCLC A549 CDX mouse model (Hu-CD34^+^-NSG-SGM3 mice) and demonstrated that bi-CD47-IT was also highly effective in the presence of a human immune system (**Fig. 4G-L** and Supplementary Fig. S4C), and with no significant changes in body weight of bi-CD47-IT treated mice relative to controls (Supplementary Fig. S4D). Immunohistochemistry (IHC) analyses using an anti-human CD47 mAb and an anti-Ki-67 mAb further confirmed that bi-CD47-IT is highly effective in significantly eliminating intra-tumor CD47 positive cells and this was associated with a significant reduction in Ki67 proliferating cells in both the orthotopic *NSG* and humanized mouse models (Supplementary Fig. S4E-H).

**Figure 4.**
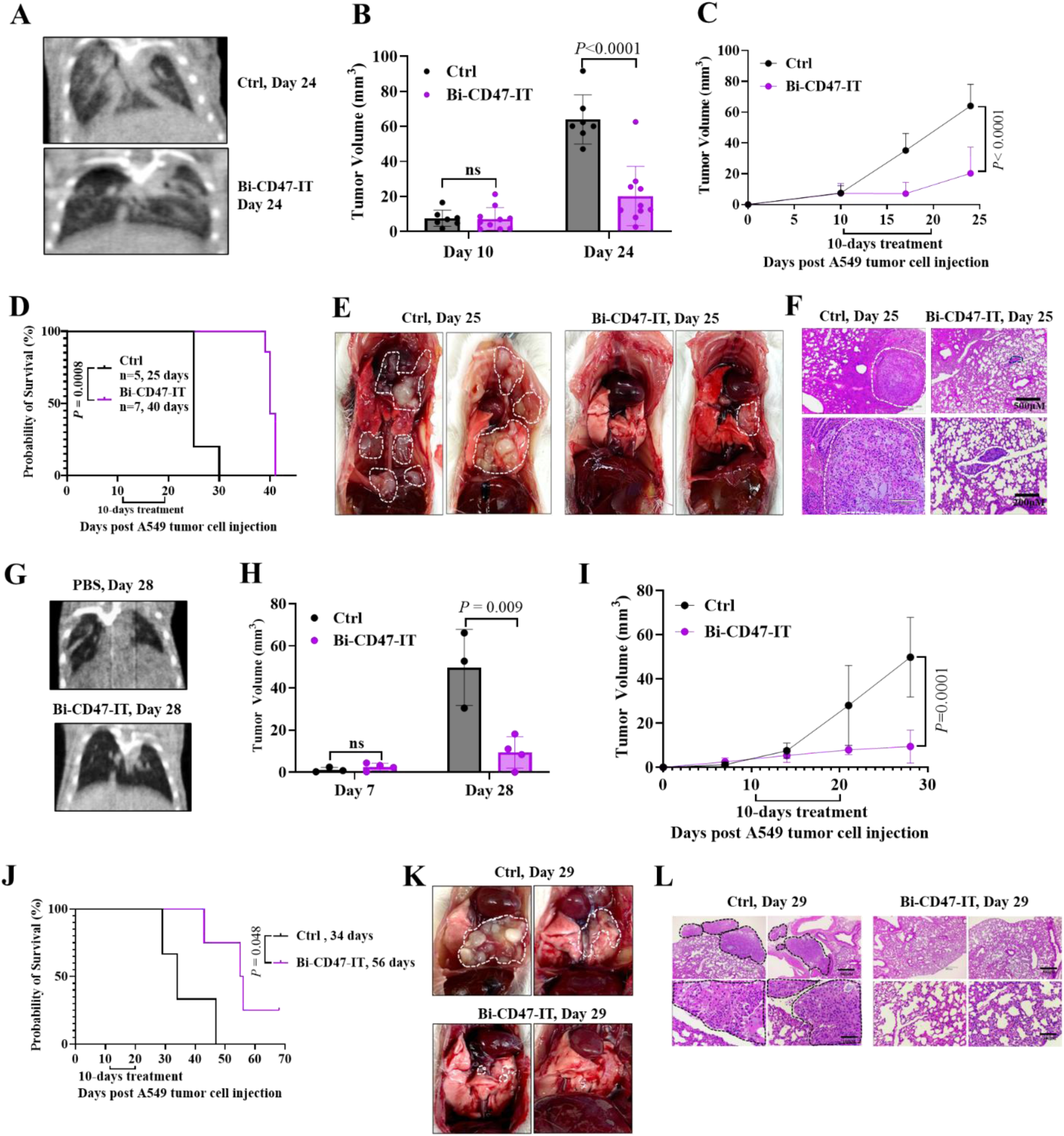
Bi-CD47-IT effectively inhibited tumor growth in orthotopic NSCLC CDX mouse models. **(A-F)** Orthotopic NSCLC A549 CDX mouse model (*NSG* mice). **(A)** Representative lungs by micro-CT scan analysis. **(B)** Tumor volumes by micro-CT scan analysis on day 10 and 24. (**C)** Tumor volume curves by micro-CT scan analysis. **(D)** Survival curves. **E)** Gross examination of the representative lungs on day 25. **(F)** Histology analysis (H&E staining) of the representative lungs on day 25. **(G-L)** Humanized orthotopic NSCLC A549 CDX mouse model (Hu-CD34^+^ NSG-SGM3 mice). **(G)** Representative lungs by micro-CT scan analysis on day 28. **(H)** Tumor volumes by micro-CT scan analysis on day 7 (pre-treatment) and 28. **(I)** Tumor volume curves by micro-CT scan analysis. **(J)** Survival curves. **(K)** Gross examination of the representative lungs on day 29. **(L)** Histology analysis (H&E staining) of the representative lungs on day 29. The *p* values in panel **(B)** and **(H)** were calculated using two-tailed Student t-test (GraphPad Prism 10). The *p* values in panel **(C)** and **(I)** were calculated using Two-way ANNOVA (GraphPad Prism 10). The *p* values in panel **(D)** and **(J)** were calculated using the Mantel-Cox log-rank test (GraphPad Prism 10).

### Bi-CD47-IT was even more effective in an experimental metastasis NSCLC A549 CDX mouse model

To test if bi-CD47-IT is effective at preventing or treating metastatic lung cancer, the major cause of death for most lung cancer patients, A549 cells (2 million) were IV injected into *NSG* mice via tail vein, and bi-CD47-IT treatment was started on day 4 (Supplementary Fig. S5A). As shown in **Fig. 5A**, bi-CD47-IT significantly prolonged the survival of the tumor-bearing mice from median survival of 36 days in controls to 75 days in the bi-CD47-IT-treated group. On day 38, two random tumor-bearing mice were euthanized from each group. Widespread tumor nodules were observed in the representative lungs, chest wall, diaphragm and abdomen of the control mice, as expected for this cell line. In contrast, no tumor nodules were observed in the representative lungs from the bi-CD47-IT-treated group, and there were no signs of metastatic lesions (**Fig. 5B**). Histological analysis (H&E staining) demonstrated large and invasive tumor masses in control mice and almost no tumor cells in the representative lungs of bi-CD47-IT-treated mice (**Fig. 5C**). The spleen was slightly enlarged in controls (**Fig. 5D**). Spleen IHC analysis using an anti-human CD47 mAb demonstrated significantly more metastatic tumor infiltration in the controls than in the bi-CD47-IT treatment group (**Fig. 5E-F**). Lung IHC analyses using an anti-human CD47 mAb and an anti-Ki-67 mAb as well as spleen IHC analysis using an anti-Ki-67 mAb further confirmed that bi-CD47-IT is highly effective in the metastatic *NSG* mouse model (Supplementary Fig. S5C–E). Gross examination and histology analysis (H&E staining) demonstrated no difference in the liver or kidney in the controls versus bi-CD47-IT-treated mice (Supplementary Fig. S5F-I). These data demonstrated that bi-CD47-IT is highly effective in the experimental metastasis NSCLC CDX mouse model, which potentially has implications for treatment of patients with advanced-stage NSCLC.

**Figure 5.**
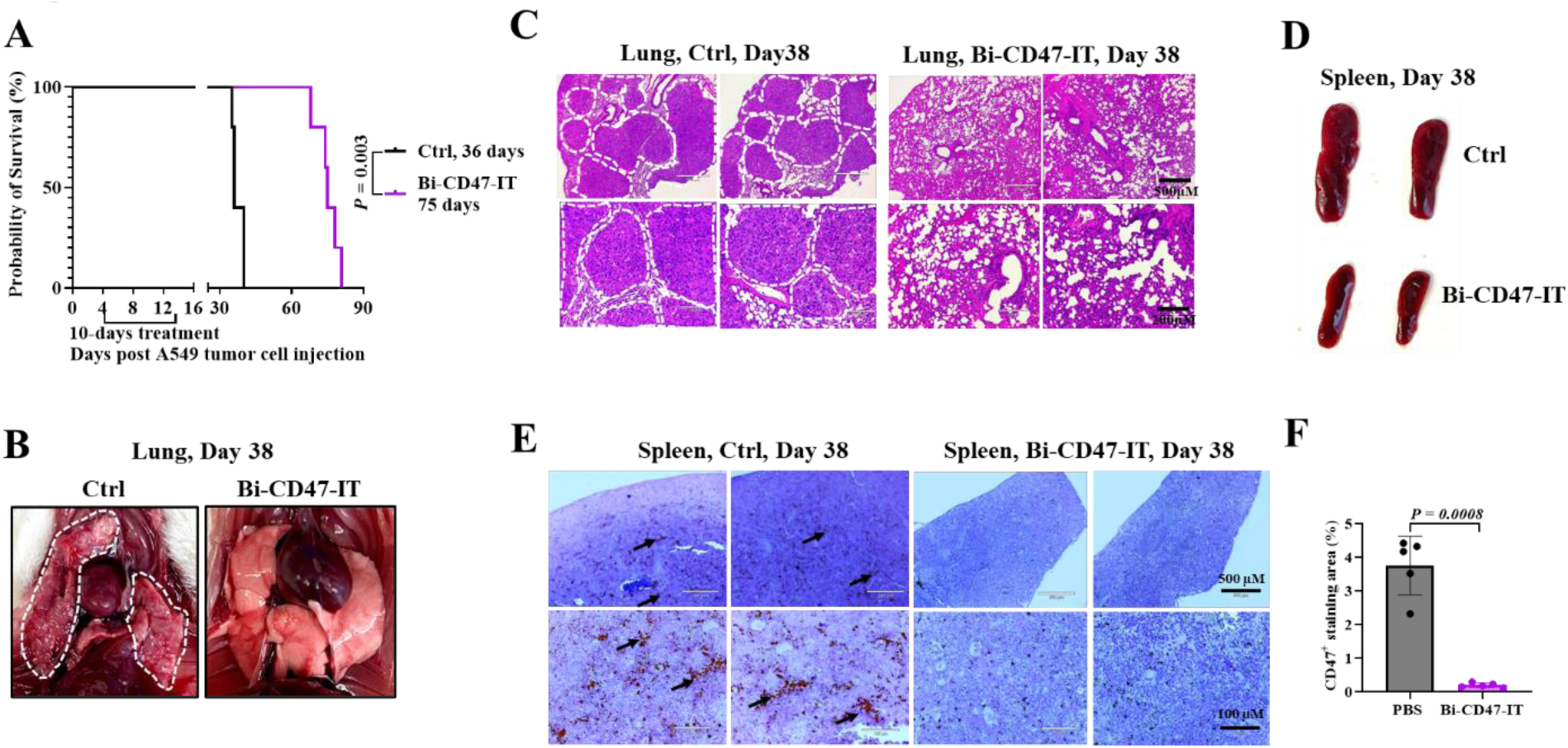
Bi-CD47-IT was even more effective in an experimental metastasis NSCLC A549 CDX mouse model. **(A)** Survival curves. **(B)** Gross examination of the representative lungs on day 38. **(C)** Histology analysis (H&E staining) of the representative lungs on day 38. **(D)** Gross examination of the representative spleens on day 38. **(E-F)** Immunohistochemistry (IHC) analysis of representative spleens using an anti-human CD47 mAb on day 38. The *p* value in panel **(A)** was calculated using the Mantel-Cox log-rank test (GraphPad Prism 10). The *p* value in panel **(F)** was calculated using two-tailed Student t-test (GraphPad Prism 10).

### Bi-CD47-IT demonstrated potent anti-tumor activity in two human NSCLC PDX NSG mouse models

Because PDXs are heterogeneous and better recapitulate lung cancer response, we further assessed the *in vivo* efficacy of bi-CD47-IT in two human NSCLC PDX *NSG* mouse models, CT-257 with a PIK3CA E542K mutation and 220900 with an EML4-ALK fusion. For PDX CT-257 and PDX 220900, bi-CD47-IT treatment was started on day 15 and day 12 post implantation, respectively (Supplemental Fig. S6A and S6C). As shown in **Fig. 6A-F**, bi-CD47-IT effectively inhibited tumor growth in both PDX models.

**Figure 6.**
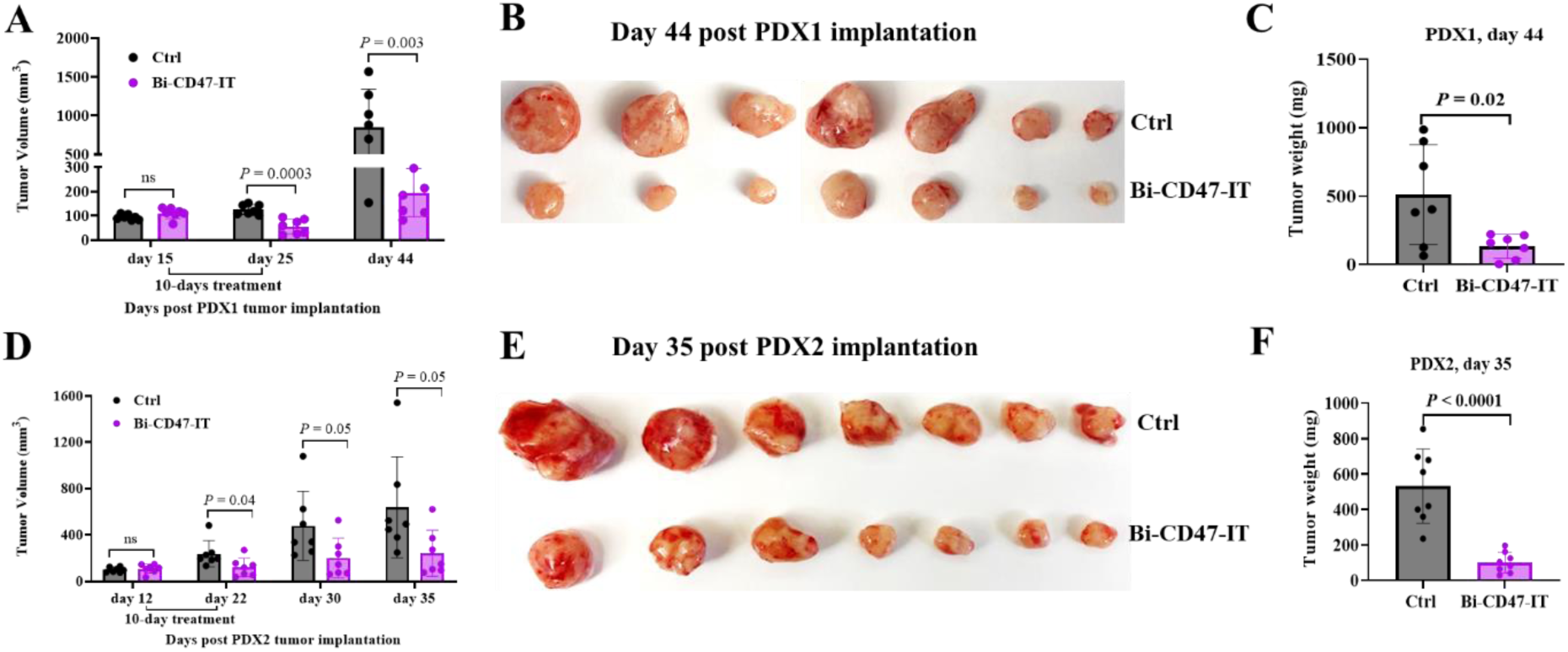
Bi-CD47-IT was highly effective in two NSCLC PDX models. (A-C) NSCLC PDX1 (CT257-F4) mouse model. **(A)** Tumor volume on day 15 (pre-treatment), 25 (post treatment) and 44 (study endpoint). **(B)** Tumor size on day 44. **(C)** Tumor weight on day 44. **(D-F)** NSCLC PDX2 (220900-M3) mouse model. **(D)** Tumor volume on day 12 (pre-treatment), 22 (post treatment), 30 and 35 (study endpoint). **(E)** Tumor size on day 35. **(F)** Tumor weight on day 35. The *p* values in panel **(A), (C), (D)** and **(F)** were calculated using two-tailed Student t-test (GraphPad Prism 10).

### Bi-CD47-IT was more effective than Cisplatin in an NSCLC A549 CDX mouse model

We hypothesized that bi-CD47-IT would be more effective in combination therapy rather than monotherapy. We chose cisplatin, a standard of care systemic chemotherapy for NSCLC. We first compared the *in vivo* efficacy of bi-CD47-IT versus cisplatin in an A549 CDX *NRG* (Rag2; IL2rg-deficient) mouse model. Treatments with bi-CD47-IT were done as detailed above and cisplatin was intraperitoneally injected at standard preclinical dose of 5 mg/kg (23,24), once weekly for 2 doses total (Supplementary Fig. S7A). As shown in Supplemental Fig. S7B-C, bi-CD47- IT as a single agent was significantly more effective than cisplatin as a single agent (*p*<0.0001), and the combination of bi-CD47-IT with cisplatin resulted in a significantly better tumor reduction than cisplatin treatment alone (*p*<0.0001). Because bi-CD47-IT was so effective, we were unable to detect a significant difference in tumor growth between the bi-CD47-IT-only versus the combination treated group, but the tumor volumes from the combination-treated group were smaller than the group treated with only bi-CD47-IT. There were no visible tumors in 2 of 6 tumor-bearing mice in the bi-CD47-IT treatment group at the study endpoint (day 28 post the tumor cell injection). Although *NRG* mice are recognized as a better model (29) for chemotherapy studies, cisplatin was still very toxic to the CDX *NRG* mice. The body weight on day 14 (post treatment) was significantly decreased in both cisplatin-only group and cisplatin + bi-CD47-IT combination treatment group (Supplementary Fig. S7D). Our data show that bi-CD47-IT is not only more effective than cisplatin, but also more tolerable in *NRG* mice.

## Discussion

CD47 is overexpressed on the surface of lung cancer cells, and on many other types of cancer cells. CD47 is also expressed on normal tissues including red blood cells and lymphocytes with relatively lower expression level. We hypothesized that CD47 receptor could be used for targeted therapy of lung cancer and other CD47^+^ cancers. We have a diphtheria toxin-based recombinant immunotoxin development platform using a unique diphtheria toxin-resistant yeast *Pichia pastoris* expression system. This yeast expression system overcomes expression and purification problems encountered with *E. coli*-based expression systems to deliver high production levels and excellent purification quality of bi-CD47-IT. We genetically engineered bi-CD47-IT for targeted therapy of lung cancer and other CD47^+^ cancers. The bi-CD47-IT demonstrated *“*optimal” binding avidity to lung cancer and other CD47^+^ cancers, no or weak binding to CD47^+^ normal tissues including human red blood cells and lymphocytes to overcome the possible toxicities to normal tissues seen with previous CD47 blockade approaches (20). For improved solid tumor penetration and better anti-tumor efficacy, our bivalent CD47 targeting moieties could facilitate internalization by CD47 receptor crosslinking, similar to the bivalent anti-CD64-IT (30). We speculate that the following features of bi-CD47-IT contributed to its high specificity: **1)** Higher binding avidity on cancer cells (due to higher CD47 expression level) than on normal cells (due to lower CD47 expression level). **2)** Shorter half-life than the parent mAb (∼40 min vs ∼17 days). **3)** Lower clinical dose than the parent mAb (<20 µg/kg vs 1-30 mg/kg). We believe that bi-CD47-IT is a promising targeted therapy drug candidate for relapsed/refractory lung cancer and other CD47^+^ cancers.

Targeted cytotoxic cancer therapy including antibody-drug conjugates and ITs have been developed (19), and ITs in particular showed remarkable efficacy in hematologic malignancies, as evidenced by the FDA approval of Lymphir^®^ (IL-2 fusion toxin) for cutaneous T-cell lymphoma (CTCL), Elzonris^®^ (IL-3 fusion toxin) for blastic plasmacytoid dendritic cell neoplasm (BPDCN). On-target off-tumor toxicity is a major concern for CD47-based therapies including bi-CD47-IT. Although we have demonstrated that bi-CD47-IT was not toxic via *in vitro* binding analysis and hemagglutination assays to human PBMC as well as *in vivo* toxicity study in humanized mice (20), currently we are continually assessing possible toxicity of bi-CD47-IT in human CD47 knock-in mice where human CD47 is expressed across all murine organs and tissues. This toxicity profile will further provide a green light for advancing the bi-CD47-IT into clinic.

We previously demonstrated that bi-CD47-IT was very compellingly effective against T-cell lymphoblastic leukemia (20). In this study, we further demonstrated that bi-CD47-IT was also effective against solid lung cancer. However, we noticed that bi-CD47-IT was relatively more effective against T-ALL than against solid lung cancer. We speculate that this is likely due to the inherent complexities of the tumor stroma and vasculature of solid tumors. The following observations in this study also support this rationale. We observed that bi-CD47-IT was more effective in the early-stage (≤50 mm^3^) than in the late-stage (>50 mm^3^) NSCLC in the subcutaneous mouse models. We also observed that bi-CD47-IT was more effective in experimental metastasis and orthotopic mouse models than in subcutaneous mouse models. Based on these observations, we speculate that combination therapy of bi-CD47-IT with other treatments such as surgical removal, cisplatin chemotherapy, radiotherapy, and targeted therapy combinations might be more effective for lung cancer treatment. Indeed, previous studies demonstrated that both cisplatin and radiotherapy enhanced CD47 expression (31,32), while the EGFR TKI, gefitinib, or the KRAS^G12C^ inhibitor, AMG 510 reduced CD47 expression (8,10), which is consistent with the oncogenic regulation of CD47 expression by EGFR and KRAS (8–10). These data provide rational therapeutic combinations to be investigated for incorporating bi-CD47-IT-targeted therapy into second and subsequent lines of therapy with standard of care lung cancer therapies.

The binding affinities of bi-CD47-IT to both A549 and H1299 cells were not high (K_D_=916.5 nM and 5,825 nM, respectively). However, the *in vitro* efficacy was very impressively high (IC_50_=1.58 × 10^-11^ M and 0.97 × 10^-10^ M, respectively) possibly due to the over expression level of CD47 on the tumor cell surface. The efficacy of the bi-CD47-IT is in accordance with a large body of preclinical evidence suggesting that targeting CD47 has excellent therapeutic potential in NSCLC, particularly in EGFR-mutant tumors. This combined with the fact that CD47 is highly expressed in large number of NSCLC emphasizes the clinical implications of bi-CD47-IT, suggesting it could be broadly applicable in lung cancer patients.

In summary, we have demonstrated that bi-CD47-IT is highly effective against NSCLC in multiple CDX and PDX mouse models including subcutaneous, orthotopic, metastatic and humanized models. Bi-CD47-IT is a novel and promising targeted therapy for NSCLC.

## Acknowledgement

This study was in part supported by the Thoracic Oncology Research Initiative (TORI) of University of Colorado Cancer Center (P30CA046934) and Colorado Office of Economic Development & International Trade Grant (DO 2025-0135).

