## Supplemental Figures 1-7 and Supplemental Table 1 for "Bivalent CD47 Immunotoxin for Targeted Therapy of Lung Cancer"

**Supplementary Data:**

**Supplementary Figures 1-7:**

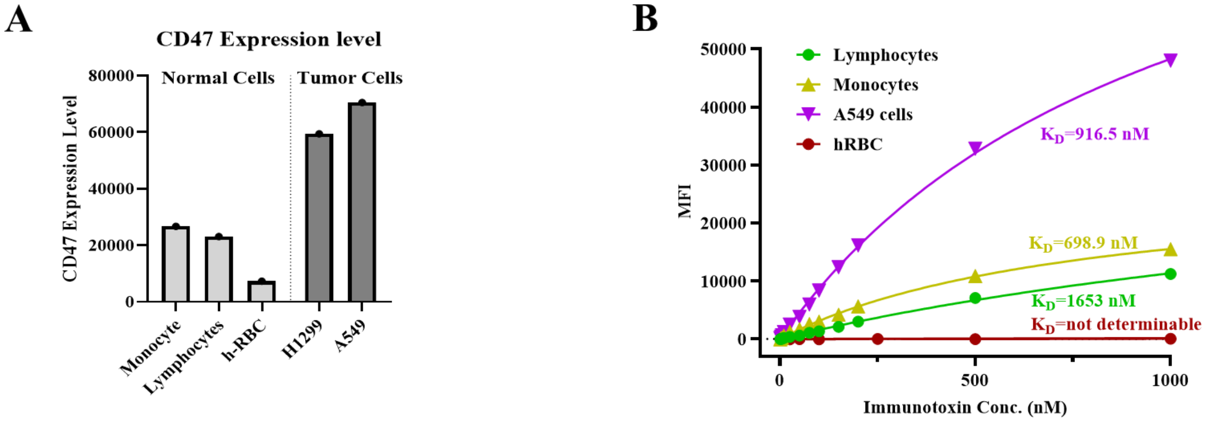


| **Sample** | **MESF** | **Ab DOL** | **CD47 Expression** |
|---|---|---|---|
| **H1299** | 300,139 | 10.1 | 59433.4 |
| **A549** | 355,888 | 10.1 | 70472.8 |
| **RBC** | 36,714 | 10.1 | 7270.1 |
| **Lymph** | 116,746 | 10.1 | 23118.1 |
| **Mono** | 134,807 | 10.1 | 26694.4 |

**Supplementary Figure 1.**

Comparison of CD47 expression levels across different cell types and the *in vitro* binding affinity of bi-CD47-IT. **(A)** CD47 expression level of human NSCLC (A549 and H1299) cells versus human red blood cells, monocytes, and lymphocytes. MESF: Molecules of equivalent soluble fluorochrome, DOL: Degree of labeling, RBC: Red blood cells. **(B)** *In vitro* binding affinity comparison of bi-CD47-IT to human NSCLC A549 cells versus human red blood cells, monocytes, and lymphocytes.

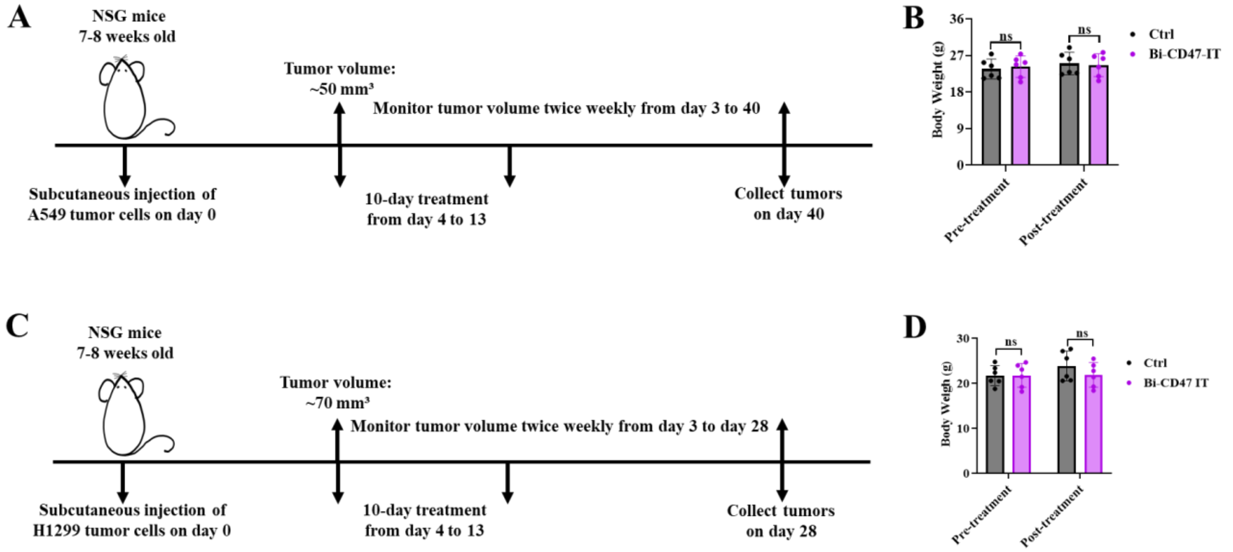


**Supplementary Figure 2.**

Bi-CD47-IT effectively inhibited tumor growth in two NSCLC CDX mouse models. **(A)** Schematic diagram of the *in vivo* efficacy study of bi-CD47-IT in a human NSCLC A549 CDX mouse model. **(B)** Body weight of the A549 tumor-bearing mice on day 3 (pre-treatment) and day 14 (post-treatment). **(C)** Schematic diagram of the *in vivo* efficacy study of bi-CD47-IT in a human NSCLC H1299 CDX mouse model. **(D)** Body weight of the H1299 tumor-bearing mice on day 3 (pre-treatment) and day 14 (post-treatment).

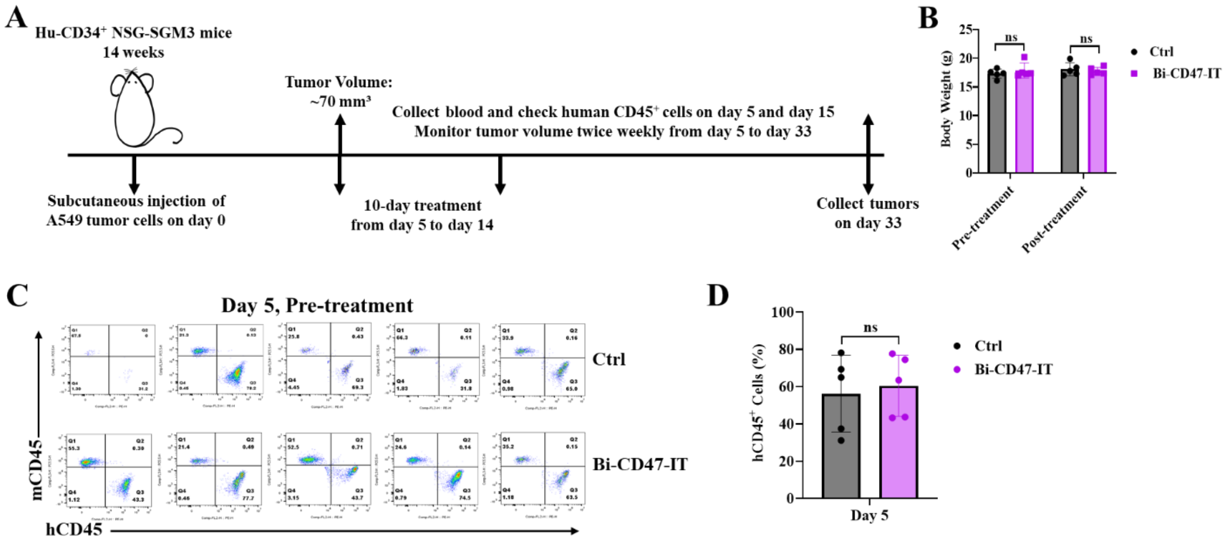


**Supplementary Figure 3.**

Bi-CD47-IT effectively inhibited tumor growth in a humanized NSCLC A549 CDX mouse model. **(A)** Schematic diagram of the *in vivo* efficacy study of bi-CD47-IT in a humanized NSCLC A549 CDX mouse model (Hu-CD34$^{+}$-NSG-SGM3 mice). **(B)** Body weight of the tumor-bearing mice on day 4 (pre-treatment) and 15 (post treatment) in the humanized NSCLC A549 CDX mouse model. **(C)** Chimerism level analysis by flow cytometry for the peripheral blood samples on day 5 (pre-treatment). **(D)** Quantified chimerism level of the peripheral blood samples on day 5 (pre-treatment). mCD45, murine CD45; hCD45, human CD45. The *p* values in panel **(B)** and **(D)** were calculated using two-tailed Student t-test (GraphPad Prism 10).

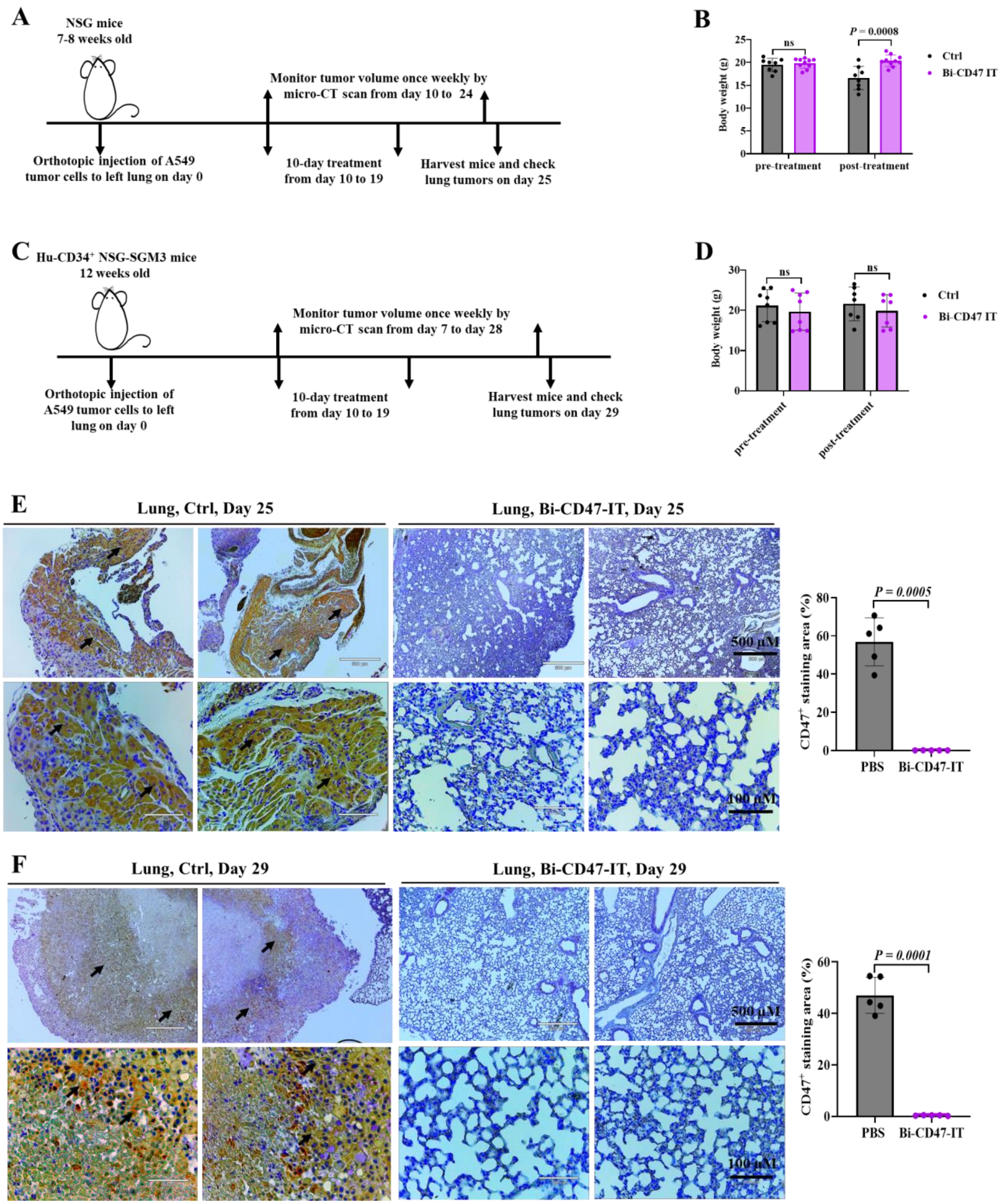
A
NSG mice
7-8 weeks old
Monitor tumor volume once weekly by micro-CT scan from day 10 to 24
Orthotopic injection of A549 tumor cells to left lung on day 0
10-day treatment from day 10 to 19
Harvest mice and check lung tumors on day 25
B
Body weight (g)
ns
P = 0.0008
pre-treatment
post-treatment
Ctrl
Bi-CD47 IT
C
Hu-CD34+ NSG-SGM3 mice
12 weeks old
Monitor tumor volume once weekly by micro-CT scan from day 7 to day 28
Orthotopic injection of A549 tumor cells to left lung on day 0
10-day treatment from day 10 to 19
Harvest mice and check lung tumors on day 29
D
Body weight (g)
ns
ns
pre-treatment
post-treatment
Ctrl
Bi-CD47 IT
E
Lung, Ctrl, Day 25
Lung, Bi-CD47-IT, Day 25
500 μM
100 μM
CD47+ staining area (%)
P = 0.0005
PBS
Bi-CD47-IT
F
Lung, Ctrl, Day 29
Lung, Bi-CD47-IT, Day 29
500 μM
100 μM
CD47+ staining area (%)
P = 0.0001
PBS
Bi-CD47-IT

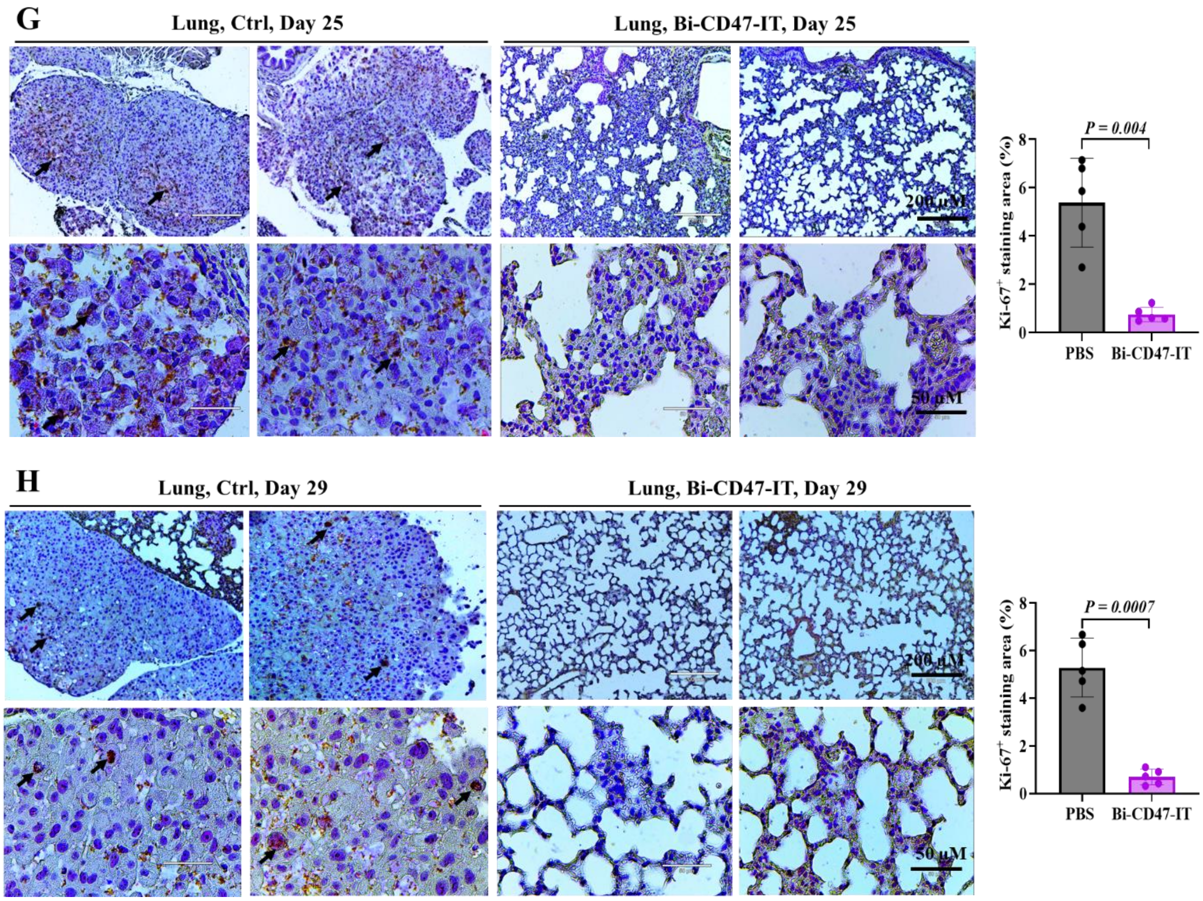


**Supplementary Figure 4.**

Bi-CD47-IT effectively inhibited tumor growth in orthotopic NSCLC CDX mouse models. **(A)** Schematic diagram of the *in vivo* efficacy study of bi-CD47-IT in an orthotopic NSCLC A549 CDX mouse model (*NSG* mice). **(B)** Body weight of the tumor-bearing mice on day 9 (pre-treatment) and day 20 (post-treatment) in the orthotopic NSCLC CDX mouse model. **(C)** Schematic diagram of the *in vivo* efficacy study of bi-CD47-IT in a humanized orthotopic NSCLC A549 CDX mouse model (Hu-CD34$^{+}$-NSG-SGM3 mice). **(D)** Body weight of the tumor-bearing mice on day 9 (pre-treatment) and day 20 (post-treatment) in the humanized orthotopic NSCLC CDX mouse model. **(E)** Immunohistochemistry (IHC) staining using an anti-human CD47 mAb on lung tissues from an orthotopic NSCLC A549 CDX *NSG* mouse model.

Brown positive staining (black arrows) was detected on most lung tissues of the vehicle control group but was absent in the lung tissues of the bi-CD47-IT treatment group. **(F)** IHC staining using an anti-human CD47 mAb on lung tissues from an orthotopic NSCLC A549 CDX humanized mouse model. Brown positive staining (black arrows) was detected on most lung tissues of the vehicle control group but was absent in the lung tissues of the bi-CD47-IT treatment group. **(G)** IHC staining using an anti-Ki-67 mAb on lung tumor cells from an orthotopic NSCLC A549 CDX *NSG* mouse model. Brown positive Ki-67 staining (black arrows) was detected in the nuclei of some tumor cells in the vehicle control group but was absent in the lung tissues of the bi-CD47-IT treatment group. This suggests that tumor cells in the control group are actively proliferating. **(H)** IHC staining using an anti-Ki-67 mAb on lung tumor cells from an orthotopic NSCLC A549 CDX humanized mouse model. Brown, positive Ki-67 staining nuclei (black arrows) were detected in some tumor cells in the vehicle control group but were absent in the lung tissues of the bi-CD47-IT treatment group. This suggests that tumor cells in the control group are actively proliferating. The *p* values in panel **(B, D and E-H)** were calculated using two-tailed Student t-test (GraphPad Prism 10).

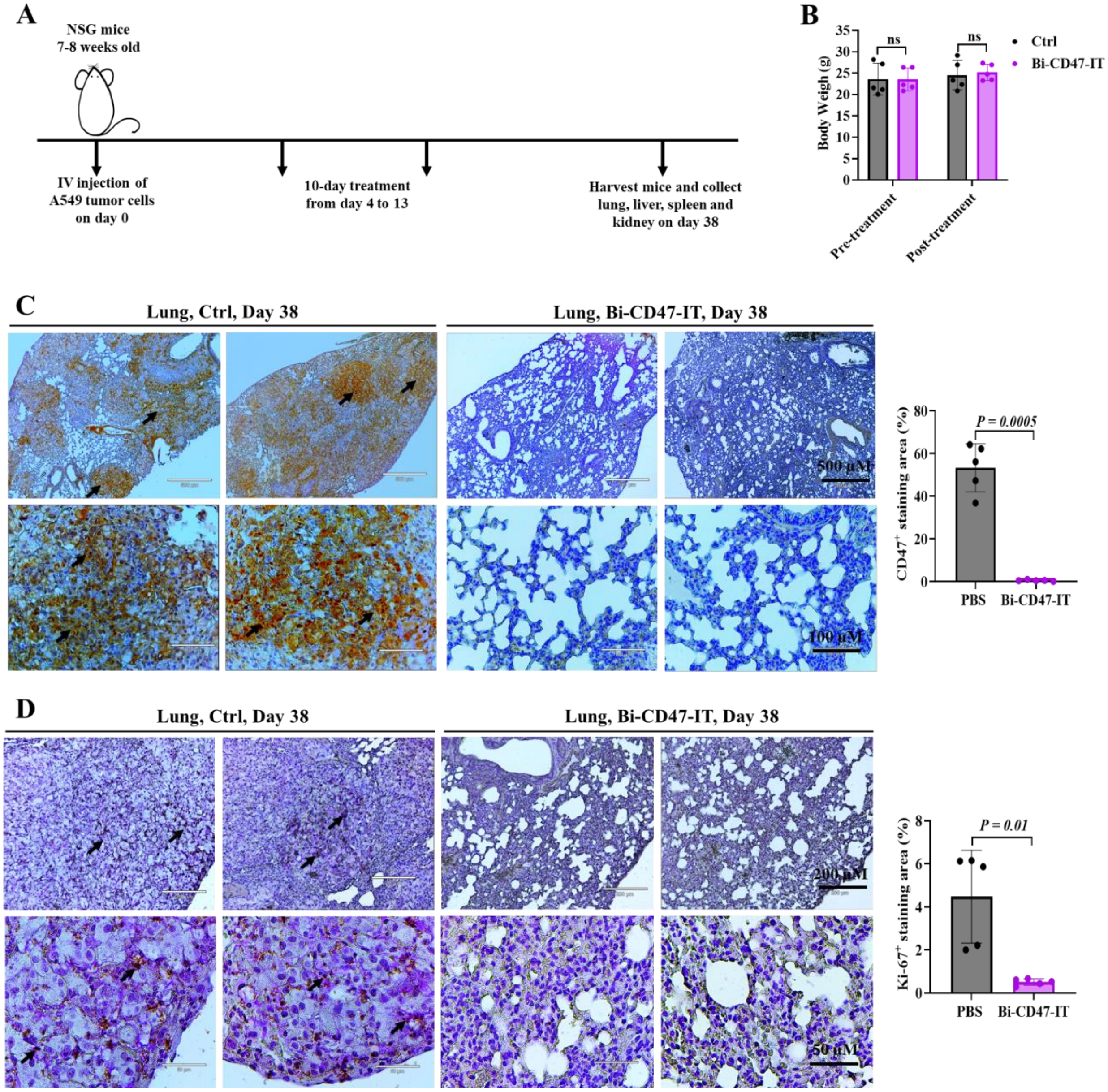
A
NSG mice
7-8 weeks old
IV injection of
A549 tumor cells
on day 0
10-day treatment
from day 4 to 13
Harvest mice and collect
lung, liver, spleen and
kidney on day 38
B
Body Weigh (g)
ns
Ctrl
Bi-CD47-IT
Pre-treatment
Post-treatment
C
Lung, Ctrl, Day 38
Lung, Bi-CD47-IT, Day 38
500 μM
100 μM
CD47+ staining area (%)
P = 0.0005
PBS
Bi-CD47-IT
D
Lung, Ctrl, Day 38
Lung, Bi-CD47-IT, Day 38
200 μM
50 μM
Ki-67+ staining area (%)
P = 0.01
PBS
Bi-CD47-IT

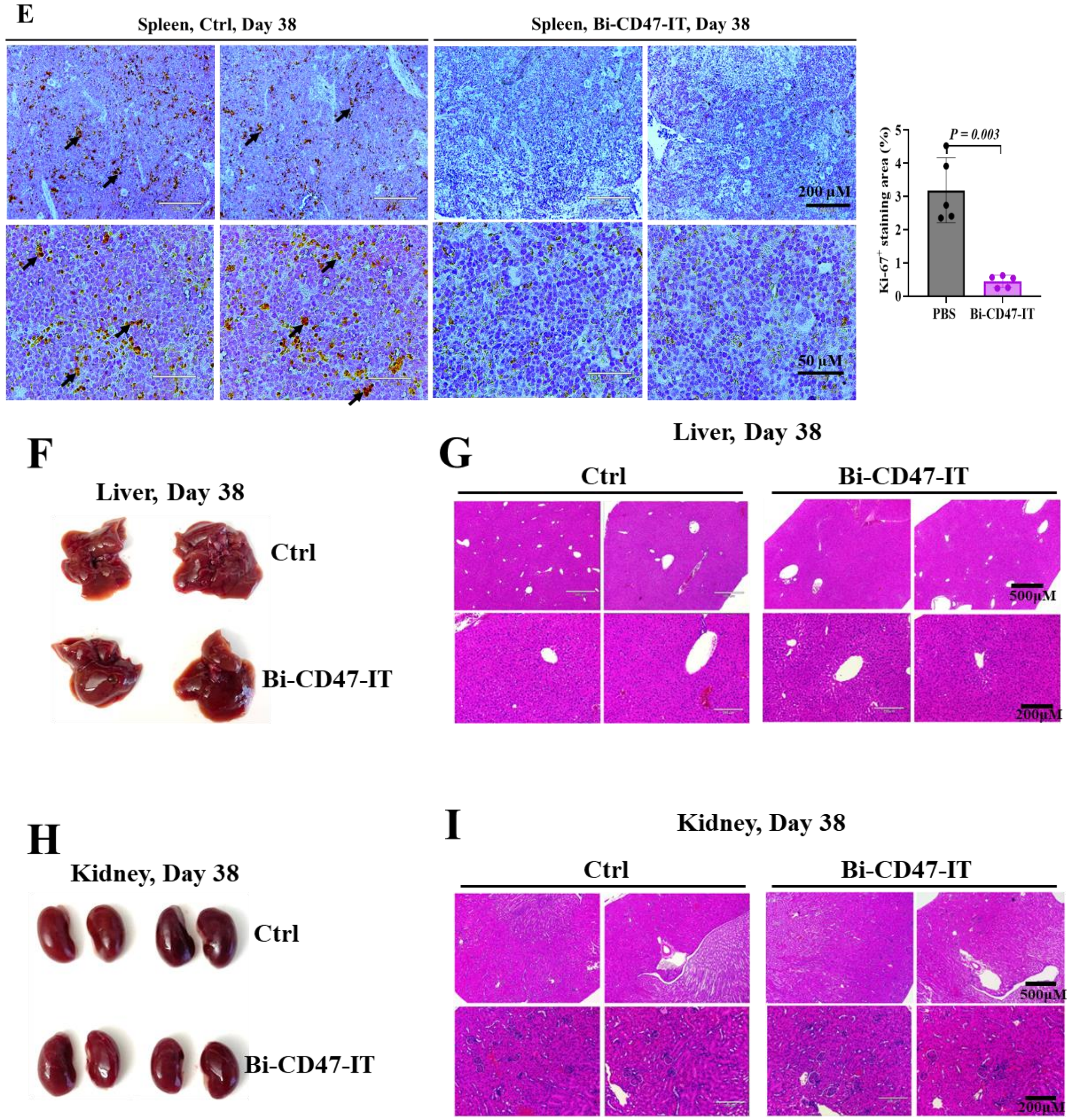


**Supplementary Figure 5.**

Bi-CD47-IT was effective in an experimental metastasis NSCLC A549 CDX mouse model. **(A)** Schematic diagram of the *in vivo* efficacy study of bi-CD47-IT in an experimental metastasis NSCLC A549 CDX *NSG* mouse model. **(B)** Body weight of the tumor-bearing mice on day 3 (pre-treatment) and day 14 (post-treatment) in the experimental metastasis model. **(C)** IHC staining using an anti-human CD47 mAb on lung tissues from a metastatic A549 NSCLC CDX *NSG* mouse model. Brown positive staining (black arrows) was detected in the vehicle control group but was absent in the bi-CD47-IT treatment group. **(D)** IHC staining using an anti-Ki-67 mAb on lung tissues from an A549 NSCLC CDX *NSG* mouse model of metastasis. Brown, positive Ki-67 staining (black arrows) was detected in the nuclei of some tumor cells in the vehicle control group but was absent in the lung tissues of the bi-CD47-IT treatment group. This suggests that tumor cells in the control group are actively proliferating. **(E)** IHC staining using an anti-Ki-67 mAb on spleens from an A549 NSCLC metastasis CDX *NSG* mouse model. Brown, positive Ki-67 staining (black arrows) was detected in some spleen-cell nuclei of the vehicle control group; however, it was absent in the spleen of the bi-CD47-IT treatment group. This suggests that the tumor cells in the control group are actively proliferating. **(F)** Gross examination of the two representative livers on day 38 post the tumor cell injection. **(G)** H&E staining of the two representative livers on day 38 post the tumor cell injection. **(H)** The gross examination of the two representative kidneys on day 38 post the tumor cell injection. **(I)** H&E staining of the two representative kidneys on day 38 post the tumor cell injection. The $p$ values in panel **(B-E)** were calculated using two-tailed Student t-test (GraphPad Prism 10).

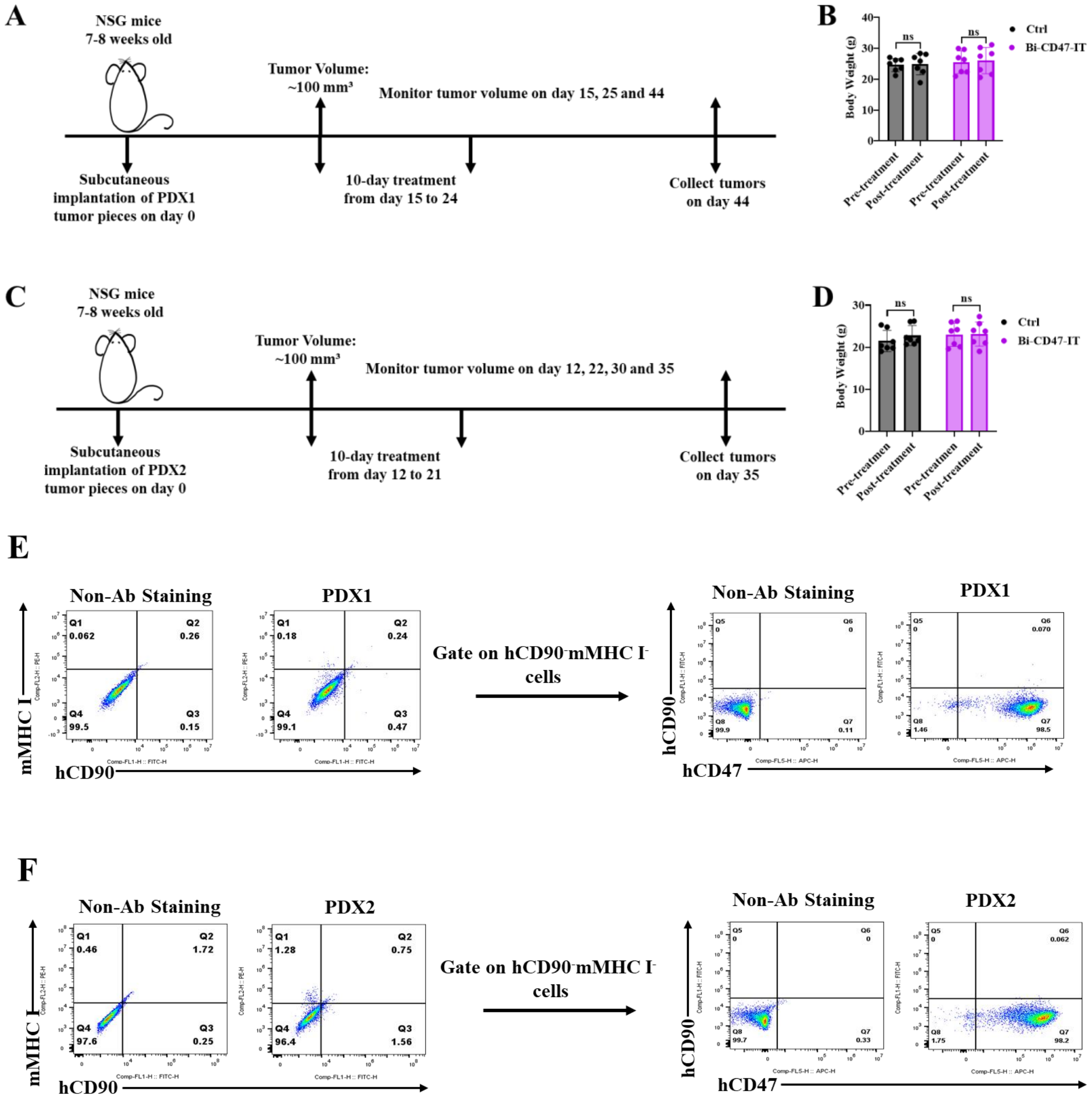


**Supplementary Figure 6.**

Bi-CD47-IT was highly effective in two NSCLC PDX models. **(A)** Schematic diagram of *in vivo* efficacy study of bi-CD47-IT in NSCLC PDX1 (CT257-F4) mouse model. **(B)** Body weight of the tumor-bearing mice on day 14 (pre-treatment) and day 25 (post-treatment) in the PDX1 mouse model. **(C)** Schematic diagram of *in vivo* efficacy study of bi-CD47-IT in the NSCLC PDX2

(220900-M3) mouse model. **(D)** Body weight of the tumor-bearing mice on day 11 (pre-treatment) and day 22 (post-treatment) in PDX2 mouse model. **(E-F)** NSCLC PDX1 and PDX2 are CD47 positive. Flow cytometry binding analysis of NSCLC PDX1 **(E)** and PDX2 **(F)** to human CD47 mAb.

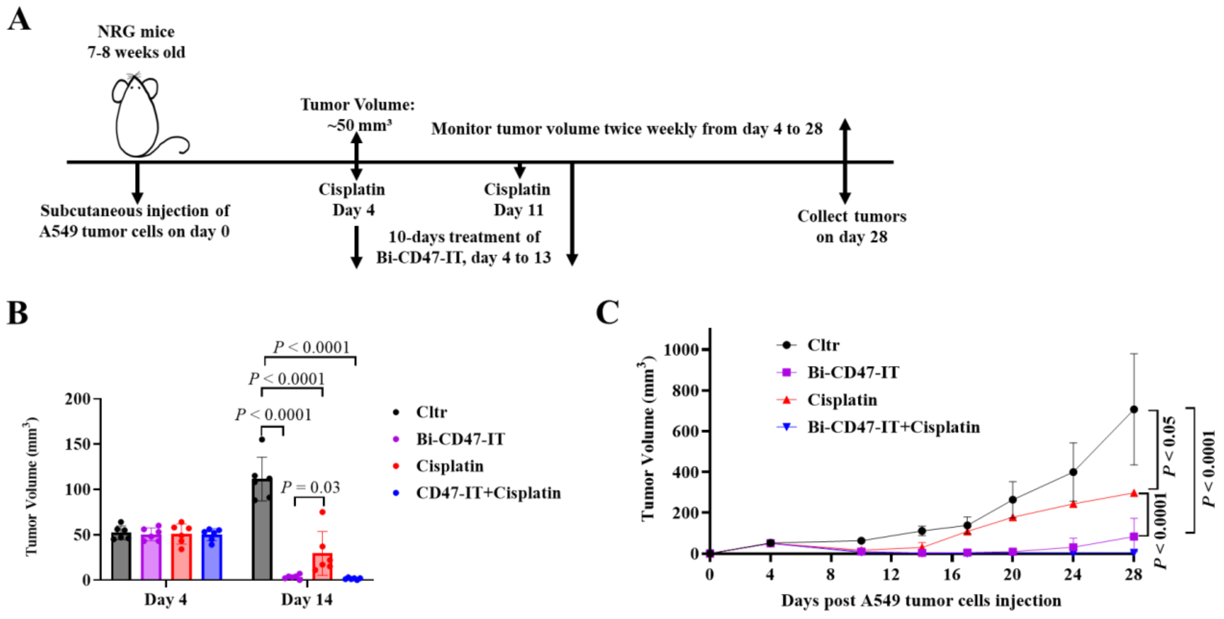


**Supplementary Figure 7.**

Bi-CD47-IT was more effective than Cisplatin in an NSCLC A549 CDX mouse model. **(A)** Schematic diagram of the *in vivo* efficacy study of bi-CD47-IT versus Cisplatin in an NSCLC A549 CDX mouse model. **(B)** Tumor volume on day 4 (pre-treatment) and 14 (post-treatment). **(C)** Tumor volume curves. **(D)** Body weight of the tumor-bearing mice on day 3 (pre-treatment) and 14 (post-treatment). The *p* values in panel **(B)** and **(D)** were calculated using two-tailed Student t-test (GraphPad Prism 10). The *p* values in panel **(C)** were calculated using Two-way ANNOVA (GraphPad Prism 10).

**Supplementary Table 1. Antibodies used in this study**

| Antibody Name | Clone | Source | Cat# |
|---|---|---|---|
| Mouse Anti-mouse MHC Class I | 2G5 | Bio-Rad | MCA2189 |
| APC Anti-human CD47 Antibody | CC2C6 | Biolegend | 323124 |
| FITC Mouse Anti-human CD47 mAb | B6H12 | BD Biosciences | 556045 |
| PerCP/Cy5.5 Anti-mouse CD45 mAb | C363-16A | Biolegend | 103314 |
| PE Anti-human CD45 Antibody | HI30 | Biolegend | 304058 |
| FITC Mouse Anti-Human CD90 | Clone 5E10 | BD Biosciences | 55595 |
| Anti-human CD47 Antibody | B6H12 | Invitrogen | 14-0479-82 |
| Anti-human Ki-67 Antibody | DVB-2 | Biocare Medical | PPM 240 DS AA |
